# Alternations of fecal microbes and lung tissue microbes in Lung Cancer patients is more consistent than that of sputum

**DOI:** 10.1101/2021.12.13.472428

**Authors:** Yanze Li, Hao Zhang, Huang Kun, Zhaohui Wang, Kai Huang

## Abstract

Microbiome in the human body environment is related to the occurrence of a variety of disease phenotypes. Recent studies discovered that lung is an open organism with in touch of air and microbe in it, and the presence of some microbes in lung cancer tissues proved that there are many microbes in lungs. In this project, we collected lung tissue, feces and sputum from three Bioprojects of NCBI related to lung cancer (LC). Each project contains LC cases and lung normal (LN) controls. Those three projects contain a total of 339 samples of 16s rRNA sequencing data. By analyzing the composition of microbes in the three environments, and predicting their functions we found that compared with sputum, the ecological environment of fecal microbe is closer to tissue microbes in terms of evolutionary relationship, indicating that the impact of feces on tissue microbes is greater than that of the sputum. We used Picrust2 to predict the differential microbe function of lung cancer (LC) and the control (Lung Normal, LN) groups in the three environments, and found that at the microbe genetic level, compared to feces and sputum, sputum and tissues, feces and tissues have more common Differential genes, at the level of differential enzyme genes and differential pathways, feces and tissues have more common differences compared to feces and sputum, sputum and tissues. Our results showed that the similarity of feces and tissue microbiome is closer than the similarity of sputum and feces microbe. Through Spearman correlation analysis based on the relative abundance of predicted pathways and the relative abundance of genus classified by LDA analysis as marker diseases and healthy samples. The results indicated that the activation of marker genus in sputum and feces and significantly changed pathways has an opposite trend, and there are many pathways contributing to glycolysis are correlated with marker genus. Patients with LC has potential to regulate the microbe composition of feces, tissues and sputum by regulating metabolism.

## Introduction

In the clinical application of microbial ecology, the previous research has proved that fecal microbe transplantation (FMT) has the potential to evaluate the patient’s abnormal and healthy phenotype[1, 2]. However, many mechanisms are currently lacking in research[3]. Current studies have shown that gut microbes can affect the host’s metabolism by producing metabolite[4] and the immune system continuously monitors resident microbiota and utilizes constitutive antimicrobial mechanisms to maintain immune homeostasis[5]. Some molecules, such as lipopolysaccharide (LPS), short chain fatty acid (SCFA), and branched chain amino acids (AAs), have been found to be involved in the occurrence of diseases[6].

Some studies have shown that in addition to the enrichment of various microbial communities in the human gastrointestinal tract[7], there also contains microbes in the oral and tissue which is in direct contact with the air environment[8]. There are also existing microbes in cancer tumors[9], and the microbes in cancer tissues are very different from those in non-tumor tissues[10, 11]. Many studies have confirmed that there are many microbes in the lung tissue[12-15]. Existing studies have shown that gut microbes can affect the composition of lung microbes through gut-lung axis, thereby affecting the development of LC [16].

Since the air way and lung tissue are connected, there are some researchers conducted that the air way microbes migrate to the lungs through air [17]. Studies have used mice to compare the influence of the environment and the intestine microbes on the lung microbes of mice and discovered the environment has more influence on the lung microbes of mice than the intestine[18]. Clinical studies have also shown that lung microbes can affected by oral microbes, lung microbes regulate lung inflammation by affecting Th17 in the lungs[19].

Studies have found that the use of metabolites in sputum as metabolic markers has contribution to the diagnosis of lung cancer[20] and microbes of feces can predict early-stage lung cancer[21]. However, whether gut microbes have an impact on the sputum and tissues, and the differences on microbe ecological function between the gut microbes, lung tissue microbes and sputum microbes is still a lack of research. In this study, we collected and analyzed 339 16s rDNA public sequencing data and found that there is a strong correlation between tissue microbes and fecal microbes. By comparing the prediction results of the microbial function of lung cancer patients and healthy people, it is found that the microbial environment of lung cancer patients’ tissues and feces have undergone more similar changes. As well as predicting the functions of microorganisms in three body environments, it was found that in the tissues, feces and sputum of patients with lung cancer, the pathways related to glycolysis metabolism have undergone significant changes, and they are widely associated with pathogenic bacteria.

## Materials and Methods

### Data

We searched for projects related to lung cancer microorganisms from the NCBI Bioproject database. Since there are few metagenomic data, we only collected projects that contained 16S rDNA sequencing data. We ended up using some of the data from the three projects. After removing the specimens not related to our subject needs, we finally collected 339 specimens, including 192 stool specimens, 97 sputum specimens, and 50 tissue specimens. The control group of tissue specimens is para-cancerous tissue. All of the runIDs from the three NCBI BioProjects are provided in Table S1 and can be downloaded from NCBI. We downloaded 16S rDNA sequencing data from the NCBI SRA database using the command-line tool fastq-dump of the SRA tools (https://github.com/ncbi/sra-tools, accessed in July 2018).

### Data processing and taxonomic assignment

We used FastQC (ver. 0.11.8; downloaded from http://www.bioinformatics.babraham.ac.uk/projects/fastqc/) to evaluate the overall quality of the downloaded data, followed by Trimmomatic (ver. 0.35; https://github.com/usadellab/Trimmomatic) to remove vector sequences and low-quality bases. We directly used single-ended sequencing reads for subsequent analyses and merged the pair-ended reads using Casper (ver. 0.8.2)[22]. We used Qiime (qiime2-2021.2) [23] to obtain the relative abundance of bacteria table. Qiime2 classifier is trained and annotated using the SILVA database. We then used Picrust2[24] to predict the KO, EC, KEGG pathways and Metacyc pathways in the microbial community.

### Statistical analysis

We uploaded all the processed data into R (ver. 4.1.0; downloaded from https://www.r-project.org). In order to remove the noise caused by low abundance bacteria, considering the bacteria in the tissue and sputum is quite a few, we removed bacteria with reads of less than 4 counts and relative abundance cutoff of 1⍰×⍰10−3. We used Picrust2 to predict the relative abundance, got the relative abundance of EC, KO and pathway, and remove the pathway data with relative abundance less than 10-6. In the LC group and the LN group, we calculated the median value of the pathway in each group, and the EC, KO and pathway of log2|Fold Change (FC)| >= 1 were defined as significant differences and used by next step analysis. The difference genus and the evolution relationship of difference genus are analyzed by LDA analysis in lefse software.

## Results

### The microbial composition of feces, tissues and sputum is quite different between LC patients and LN

The feces, tissue and sputum flora of LC patients are significantly different from those of healthy people. Using PCoA analysis based on the Bray Curtis distance algorithm, we found that feces, tissue and sputum, the LC group and the LN group can be distinguished significantly (Fig1. A). It might indicate that in the relative abundance of bacteria, sputum, lung tissue and feces are significant differences between the healthy group and the control group. Among the consistent bacteria of these three groups, the microbiome in feces, sputum and tissues have different relative abundance (Fig1. B). The boxplot indicated that the overall trends of fecal microbe and sputum microbe are more resemblance, while the evolutionary tree also shows that bacteria enriched in feces, tissues and sputum have evolutionary similarities (Fig S1). The tissue microbe is very different from these of the feces and sputum (Fig1. B), indicating that most bacteria do not migrate directly.

**Figure 1.**
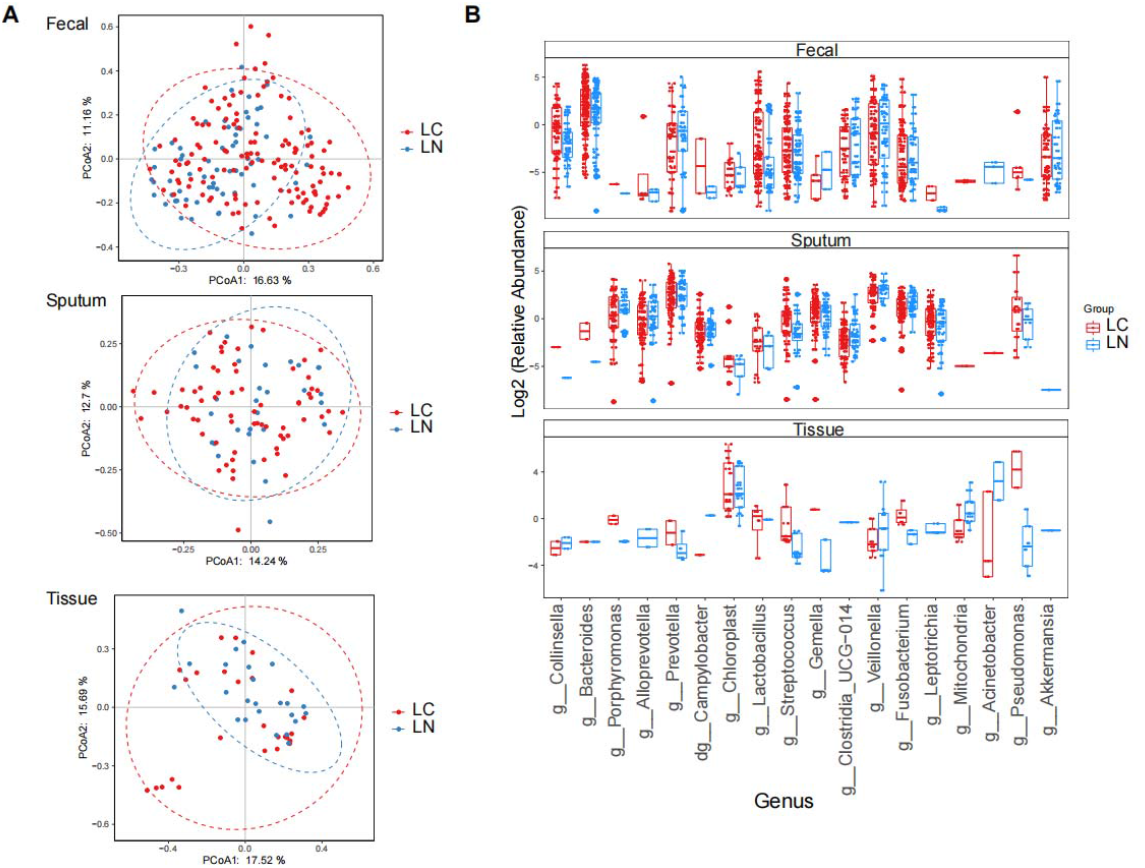
There are differences between tissues, sputum and fecal microbes, and there are also microbes that co-exist. (A) Principal coordinate analysis (PCoA) based on Bray-Curtis distance at genus level of feces, sputum and tissue showed that the overall microbiota composition was different between LC and LN. (B) The 18 genus co-existing in feces, tissues and sputum showed many differences on relative abundance.

### Compared to the sputum microbiome, the lung tissue microbiome is more consistent with the fecal microbiome

Cluster analysis is performed on the three sets of data based on the weighted unifrac emperor algorithm (Fig2. A). The results showed that the composition of fecal microbiome and sputum microbiome has a longer distance. Using the Permutational multivariate analysis of variance (Permanova) distance to visualize the distance between the three groups of samples (Fig2. B, C and D), it can be seen apparently that the distance between sputum, tissue, and feces. The closest distance to tissue is feces (Fig2. D), and the closest distance to feces is sputum (Fig2.C and E). The microbiome of feces has closest distance to sputum and tissue (Fig2. C and D). Those results showed a vital importance of fecal microbiome to tissue.

**Figure 2.**
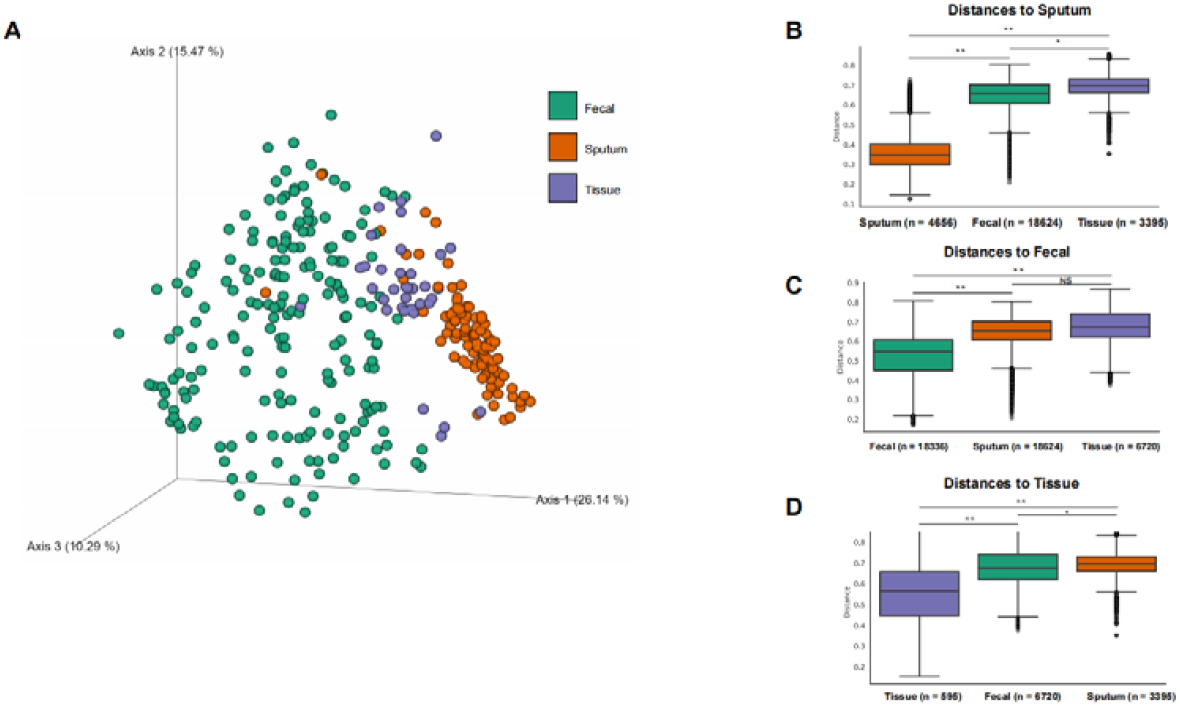
The overall composition of fecal microbes and tissue microbes has the highest similarity. (A) Weighted unifrac emperor algorithm clusters the evolutionary relationship of the 16S data of bacteria, showed that the closest evolutionary relationship to the fecal microbe is the tissue microbe. (B) Permutational multivariate analysis of variance (Permanova) distance showed that the closest distance to sputum microbe is feces, and the next closest is tissue. (C) Box plot showed that the distance to fecal microbe with sputum and tissue didn’t have significant different. (D) Box plot showed that the distance to tissue microbe is feces, and the next closest is sputum. Level of significance: **, P < 0.01; *, P < 0.05; NS, P ≥ 0.05.

### The relative abundance of bacteria can distinguish most LCs from LNs samples

In order to evaluate the different distinguishing ability of these body sets of data for LC, and explore the phenomenon causes the differences in classification ability. We used the random forest model to determine the most influential position of microbiome between LC and LN patient (Fig3. A).

**Figure 3.**
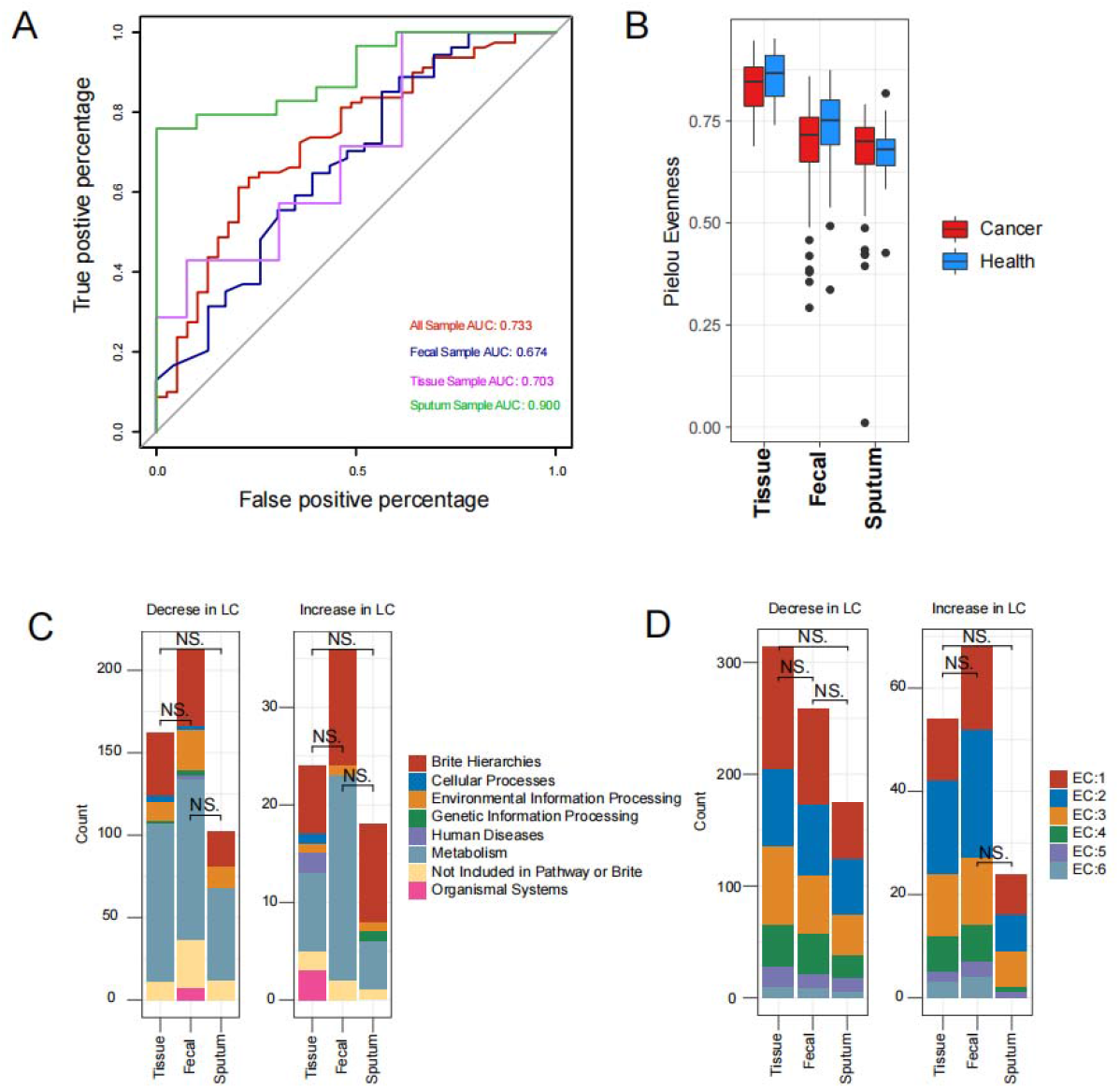
Relative abundance can recognized feces tissue and sputum between LC and LN, and the KO and EC have consistent differences on composition. (A) The Random Forest model classifies bacteria with relative abundance> 0.01%, and the results show that the relative abundance of bacteria can distinguish between the LC group and the LN group. (B) Analysis on the α diversity showed in LC group the diversity of tissues and feces decrease, and the diversity of sputum increases. (C) There was no significant difference (|log2 FC| >=0) in categorical composition between the increased and decreased KOs in the LC group. (D) There was no significant difference (|log2 FC| >=0) in the classification composition between the elevated and lowered ECs in the LC group. NS. Level of significance: P ≥ 0.05, using two-tailed t-test.

We used the random forest model to determine the characterization of microorganisms in three environments. Defining the AUC greater than 0.5 indicates that the difference between the two groups of LC and LN samples is significant. The AUC of all 339 specimens is 0.733, the AUC of fecal samples is 0.674, and the AUC of tissue samples is 0.703 (Fig3. A). The highest classification result is sputum, with an AUC of 0.900, which is consistent with the previous study [25] (Fig3. A). Most of the top 15 genus that affect the classification effect are unrecognized bacteria (Fig S2).

In order to find out the reason that RF model classification effect of sputum is the best, we did analysis on the α diversity of microbiome in these three environments (Fig3. B). The results of the diversity analysis showed that sputum had lower diversity than tissue and fecal samples (Fig3. B). Due to the lower microbial α diversity, there are fewer significant bacteria related to the occurrence and development of LCs. We then used *Picrust2* to predict the relative abundance of enzyme genes in these samples, and calculated the count of each EC (Enzyme Commission’s classification) type of enzymes with significant differences (|log2 FC| > 1) between LCs and control LNs (Fig3. C and D). We calculated the ECs of three groups using two-tailed t-test, and found that there is no significant difference in composition classification between EC and KO in the three environments (Fig3. C and D). Among them, *Veillonella* in feces. *Streptococcus* in sputum and *Streptococcus* and *Veillonella* in tissues were also found in the lower airways of patients with lung cancer. Those genera were enriched for oral taxa, which was associated with up-regulation of the ERK and PI3K signaling pathways [26].

### Significant different KO, EC and pathways in feces and tissues are more resemblance

To explore the relationship between bacterial differences and functional differences between the LC and LN groups, we used *picrust2* to predict the microbial community’s functional composition, and map the significant differences (|log2FC| >= 1) between the LC and LN groups. The Venn plot showed the count and intersections of significant different ECs and KOs (KEGG Orthology) between tissues, feces and sputum (Figure4. A and B). The genes and enzymes that have undergone significant changes in the three environments, and the feces and tissues have higher Jaccard similarity in enzymes and genes (Figure4. A). Then we plot a heatmap on the relative abundances of the two specific genes in these parts, and found that most of the common differential enzymes in feces and sputum does not exist in tissue (40/21; 52.5%), and the rest of have no relative abundance changes (FigureS3. A and B). Most of the differential enzyme genes in feces and tissues are also present in sputum (64/43; 67.2%), others have no significant change in sputum (Figure4. C). The relative abundance changes revealed the functional differences between tissues and fecal microbiome are closer than that of feces and sputum. The main common differential enzymes in feces and tissues are EC1, while the main differential enzyme type between feces and sputum is EC2, as well as tissue and sputum (Figure4. C and Figure S3).

**Figure 4.**
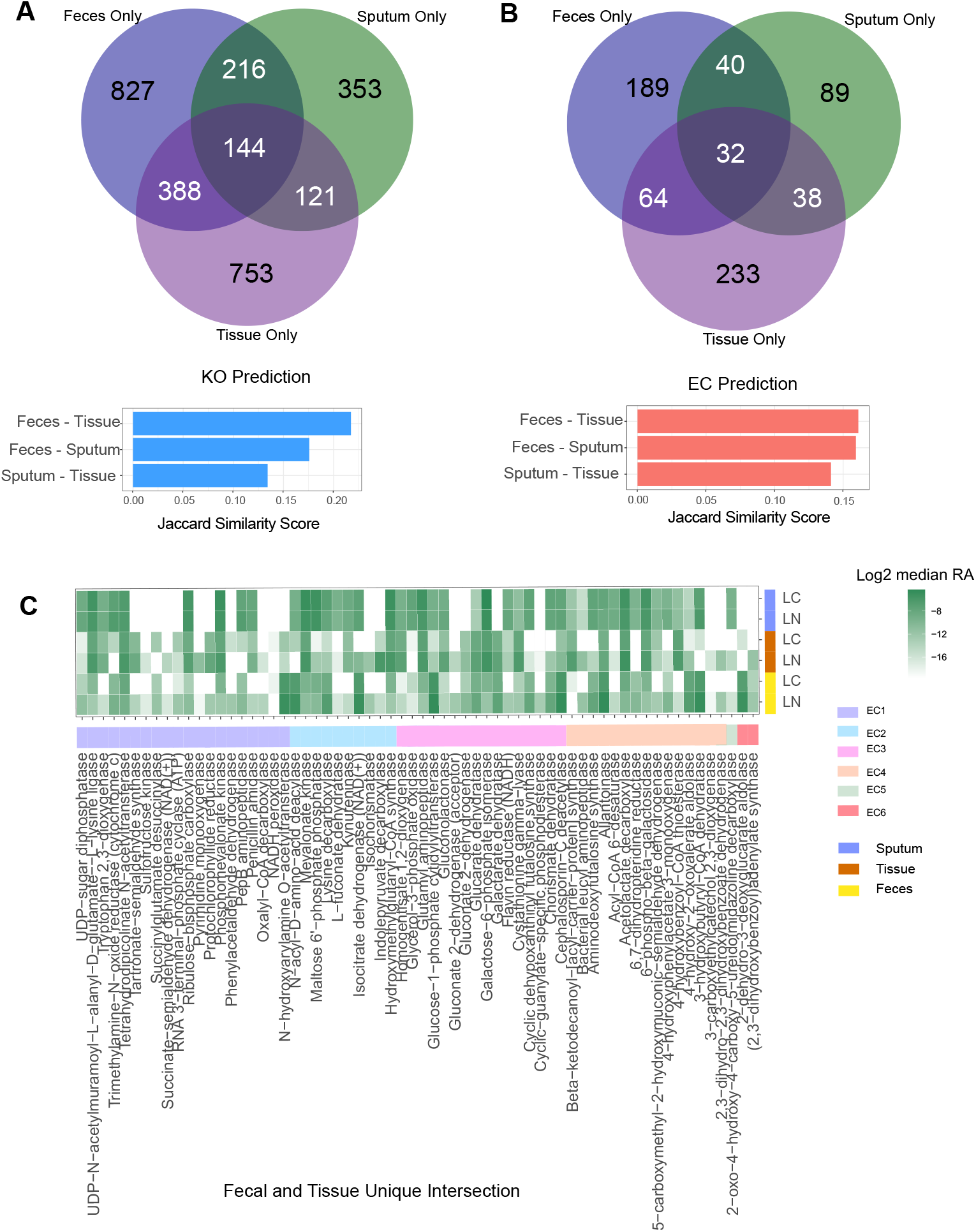
Fecal and tissue microbiome have more similarity on KO and EC. (A) Among the three sets of data, the KO that has undergone significant changes has the highest Jaccard Similarity in tissue and feces. (B) Among the three sets of data, the EC that has undergone significant changes has the highest Jaccard Similarity in tissue and feces. (C) At the EC level, most of (67.2%) the significantly different enzyme genes present in tissues and feces are also present in sputum, but there is no significant change.

### The metabolic pattern of sputum microbe in LC patients is significantly different from that of feces and tissue

We used the LDA score (P<0.05) of the lefse software to screen out the genus that had significant changes in the three environments (Fig. S4). And by comparing the KEGG database and the Metacyc database, it is found that these marker genus and significantly different pathways have different relationships in different environments. The heat map shows that the sputum marker genus has significant differences in metabolic pathways (Fig. 5A and S5). The bacteria related pathway enriched in the feces of healthy people is positively correlated with the D−galactarate degradation I, D−glucarate degradation I and superpathway of D−glucarate and D−galactarate degradation (Fig5. B, D, E and FigS5), while the pathway of sputum in healthy people is negatively correlated (Fig5. B, D, E and FigS5). The bacteria related pathway enriched in sputum and feces of LCs is positively correlated with lactose and galactose degradation I pathway, while the pathway of tissue in LCs is negatively correlated (Figure5 C). Indicating in intestinal microbes, the relative abundance of metabolic pathway using glucose in the in LC patients has decreased, while the glucose metabolism pathways of sputum have increased (Fig5. B, D, E), and the degradation activities of LCs also increased (Figure S5). This might explain why the sputum of LC patients has a higher microbial diversity, because of increase of glycan and degradation activity in the microbe of LC sputum. The reduction of biosynthetic activities and the increase of organic substances have made organic substances in the sputum environment more complicated [20]. Through enrichment analysis of the differential pathways and differential microbe in the three environments, we found that the microbe involved in monosaccharide metabolism decreased in the patient’s feces and increased in the sputum (Figure S5). The pathways related to glucose metabolism have the most extensive correlation with the marker genus that causes LC (Figure S6). These results indicating that the glycolysis metabolic feature has significantly changed between LCs and LNs.

**Figure 5.**
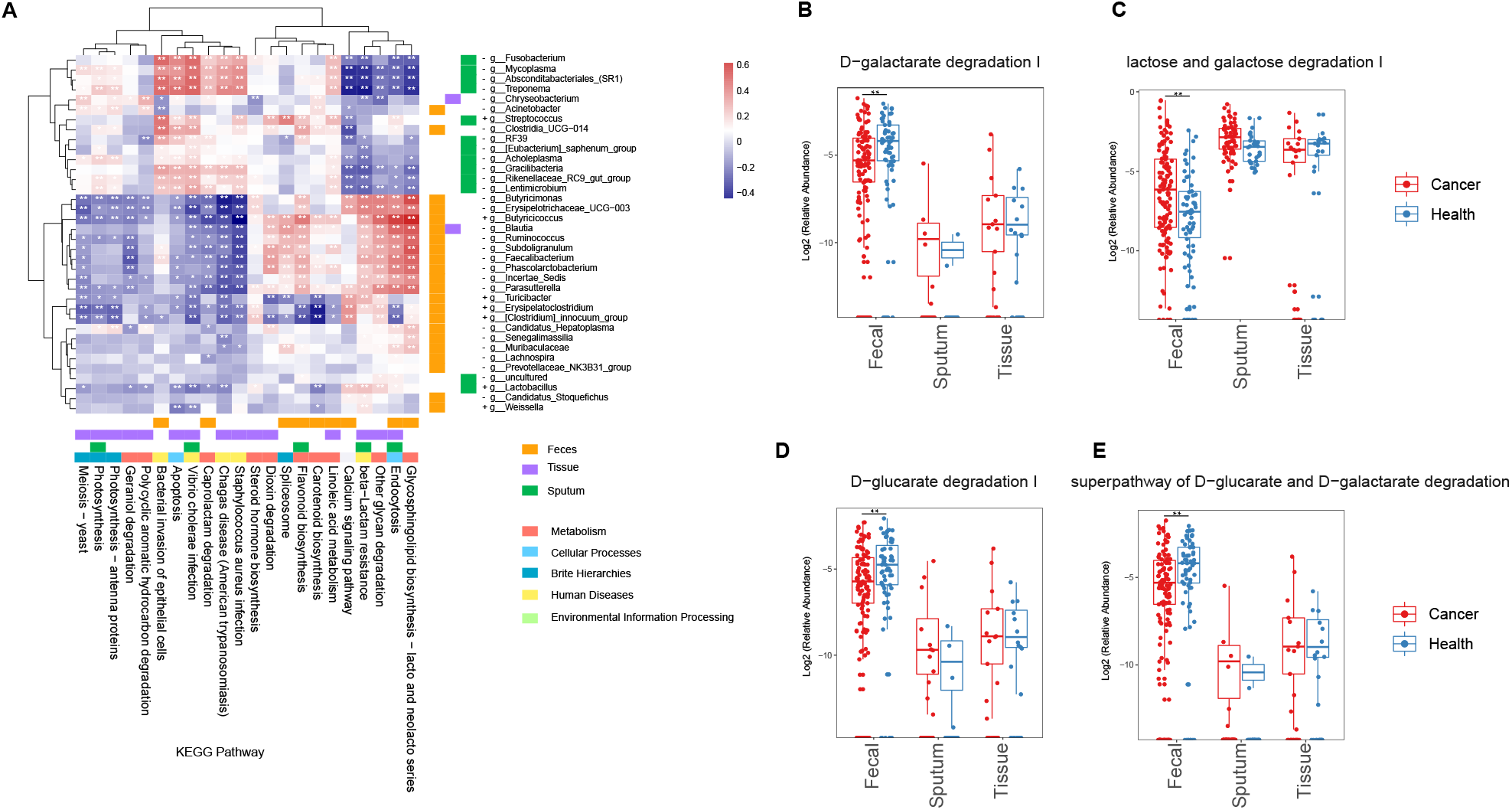
Significant changed metabolic pathways are caused by significant changes genera, and there are differences in glucose metabolism pathways between the LC and LN groups. (A) The genus that has undergone a significant change caused a significant change in the pathway. It can be seen that feces and tissues are closer in the pathway, and sputum is significantly different from feces and tissues. The green bar represents sputum, the purple bar represents tissue, and the yellow bar represents stool. The +, marked before genus means enrichment in the LC group, and -, means enrichment in the LN group. (B) The difference between D−galactarate degradation I pathway in tissue, LC of feces and sputum, LN group, feces and sputum have opposite trends. (C) The difference between lactose and galactose degradation I pathway in tissue, LC of feces and sputum, LN group, feces and sputum have same trends. (D) The difference between D−galactarate degradation I pathway in tissue, LC of feces and sputum, LN group, feces and sputum have opposite trends. (E) The difference between D−glucarate and D−galactarate degradation pathway in tissue, LC of feces and sputum, LN group, feces and sputum have opposite trends. Level of significance: **, P < 0.01; *, P < 0.05; NS, P ≥ 0.05.

## Discussion

The microbial composition of the lung tissue is related to the malignant diseases of the lungs [8]. The respiratory tract, as an open connection environment with the lungs, exchanges flora with lung microorganisms, and is also related to the occurrence of lung disease [27]. We evaluated the tissues, feces and sputum specimens of 339 LC and LN, and found that feces and tissues had a higher similarity in bacterial composition and functional composition than sputum [Fig2 and Fig4].

In this work, we found that the microbes in these three parts are different between the lung cancer and healthy groups, and there are 18 genera existing in all these three parts [Fig.1]. We used a Random Forest-based machine learning model to verify that the relative abundance of microbes can distinguish specimens of healthy people with lung cancer [Fig3]. We used LDA score to analyze the marker genus in the microbial sampling in the three human environments, and obtained the genus enriched in the LC and NC groups in those three environments [Fig. S4]. We used *Picrust2* to predict the function of the microbial community, by analyzing the three groups of microbial environments, we found that the KO and EC that have significantly changed in tissues and feces have the highest Jaccard Similarity.

Through Spearman correlation analysis of the relative abundance of differential pathways and differential genes, the correlation between the three groups of differential genus and differential pathways was demonstrated [Fig5. And Fig S5]. The genus enriched in the sputum of healthy people participates in the regulation of pathways and is enriched in tissues. The genus of the set is significantly different, and the pathways involved in glucose metabolism in feces and sputum in the LC and LN groups have a significant negative correlation. It may be related to the Warburg effect of tumor cells[28]. We assumed that the metabolism of tissue microorganisms has the potential to be interfered by fecal microbe.

The microbes in the tissues, sputum and feces of patients with lung cancer are very different from those of healthy people, and there are certain changes from composition to function. Through our analysis, we found that the microbial composition in the tissues, sputum and feces of lung cancer patients can distinguish lung cancer from healthy people. The changes in these microorganisms have resulted in changes in the genes encoding proteins, enzymes, and corresponding pathways in the genome of environmental microbes. The changes in tissue and stool are the most similar, and the functional changes between stool and sputum, tissue and sputum are very different. It is suggested that the changes in the characteristics of microbial metabolism in patients with lung cancer, especially the changes in the related characteristics of glucose metabolism, may be the cause of the microbial differences in tissues, feces and sputum.

## Author Contributions

YL and KH conceived and designed the study. YL and HZ analyzed the data. YL wrote the draft, KH and ZW revised it. All authors approved the final version.

